# A Learning-Based Formulation of Parametric Curve Fitting for Bioimage Analysis

**DOI:** 10.1101/2020.01.10.901702

**Authors:** Soham Mandal, Virginie Uhlmann

**Affiliations:** European Bioinformatics Institute (EMBL-EBI), European Molecular Biology Laboratory, Cambridge, UK

## Abstract

Parametric curve models are convenient to describe and quantitatively characterize the contour of objects in bioimages. Unfortunately, designing algorithms to fit smoothly such models onto image data classically requires significant domain expertise. Here, we propose a convolutional neural network-based approach to predict a continuous parametric representation of the outline of biological objects. We successfully apply our method on the Kaggle 2018 Data Science Bowl dataset composed of a varied collection of images of cell nuclei. This work is a first step towards user-friendly bioimage analysis tools that extract continuously-defined representations of objects.

## 1 Introduction

Parametric curve models have been extensively used in the past in the context of active contours for image segmentation [1]. In its classical formulation, a parametric active contour algorithm requires the definition of a curve model, which is then initialized in the image and evolves to capture the boundaries of an object of interest by minimizing a handcrafted cost functional referred to as the energy [2]. In practice, designing an energy implies formalizing visual intuition in mathematical terms and requires expert domain knowledge [3], limiting the usability of parametric active contour methods. The energy most often consists in a combination of multiple terms, which must be weighted properly and are difficult to robustly optimize at once. In the context of bioimage segmentation, parametric active contours have mostly fallen into disuse with the advent of more efficient, robust, and generalizable learning-based methods such as convolution neural networks [4] and random forests [5]. Being able to model the outline of objects in a (possibly already segmented) image as parametric curves however remains of interest. The crux of bioimage analysis indeed consists of extracting quantitative measurements to describe, model and understand living phenomena [6]. Continuously-defined parametric curve models offer a convenient representation to extract morphological descriptors without discretization artefacts. Discrete segmentation masks are in fact often continuously interpolated prior to quantitative analysis [7]. Devising the best set of (discrete) contour points to interpolate is, however, not trivial, as mask boundaries are likely to be noisy. It thus appears that parametric curve fitting, as historically considered in active contour methods, deserves to be considered and modernized as a tool to retrieve optimal continuously-defined representations from discrete sets of connected pixels. In this work, we present a first attempt in this direction: we rely on a convolutional neural network (CNN) to predict the best parametric curve fit onto an object contour. In that way, we trade the energy design step, which requires domain-expert knowledge, for a data-driven approach. Our network is trained with a loss designed to penalise the discrepancy between a discrete ground truth mask and a sampled version of the predicted continuous contour. From this, it learns to generate the continuous representation that most accurately approximates the discrete contour of an object.

Using neural networks to predict a set of interpolation points and retrieve a continuous model of the contour of objects in images has recently attracted attention. Most relevant to our work are those of [8] and [9]. In [8], a joint structure composed of a CNN and an autoencoder is proposed to smoothly model the surface between vertebral bodies and posterior elements in the vertebra. The surface model is a thin-plate spline, whose control points are predicted using a shape model of vertebral bodies. While yielding very promising results, this approach heavily relies on the structural specificity of the vertebra and therefore has limited application in bioimages featuring other objects. The more general problem of predicting the locations of the set of control point of a parametric spline curve that best represents an object’s contour is explored in [9]. The task is formulated as deducing the values of a variable length sequence of coefficients. The proposed solution consists of the combination of a CNN and a recurrent neural network. The loss is defined from a set of ideal, ground truth control point locations. Such a construction assumes that a unique set of control points yields the best parametric curve representation of an object contour. Since the parametrization of a continuously-defined contour is not unique, several control point sets can generate the same curve. The considered loss thus imposes artificial restrictions on the contour representation. Here, we alleviate this by adopting a curve-based (instead of control-points-based) loss.

In Section 2, we recall the formulation of a parametric curve model built using spline interpolation, present the architecture of our network, and discuss the design of the loss, which is central to our work. Section 3 is devoted to experiments: we tested our network on the Kaggle 2018 Data Science Bowl dataset [10], which contains images of cell nuclei acquired under a variety of conditions, and that vary in the cell type, magnification, and imaging modality. Finally, in Section 4, we conclude with a brief discussion and explore future directions.

## 2 Description of the Method

### 2.1 Parametric Curve Model

Our parametric curve model is constructed relying on uniform spline interpolation as in [2]. In the 2D plane, a closed parametric spline curve **r**(*t*), 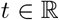 is expressed by two coordinate functions *x*_1_(*t*) and *x*_2_(*t*) as

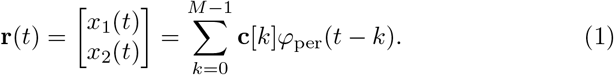

The coefficients 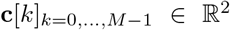 are the control points and, in our use-case, correspond to coordinates in the image space. The number of control points, *M* relate to the flexibility of the curve, as smaller number of control points yield more rigid contours. In our case, the function 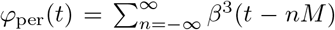 is the *M*-periodized version of the cubic B-spline generator [11]. We rely on cubic B-splines for their good approximation properties of smooth curves [12]. Our method would however equivalently work with other interpolation kernels 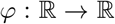, such as exponential splines [13].

### 2.2 Neural Network Architecture

Using a neural network, we aim at robustly determining how control points should be distributed along the boundary of objects in a variety of bioimages. CNN are the tools of choice when it comes to understanding spatial information from image content as they explicitly connect neurons that are spatially near in consecutive layers [14]. In particular, CNN already achieve state-of-the-art performance in several image analysis tasks such as image classification, image segmentation, and bounding box detection.

We adopt the architecture depicted in Figure 1. This construction, consisting of blocks of convolutional layers followed by pooling and ending with a dense fully connected layer, is typical of a classical CNN [15]. For a two-dimensional parametric spline curve composed of *M* control points, the final layer of our architecture is composed of 2M nodes, where each subsequent pair of nodes correspond to the image coordinates of a control point. The network architecture, as well as hyperparameters such as learning rate and batch size, could be adapted and optimized for specific biological objects.

**Fig. 1:**
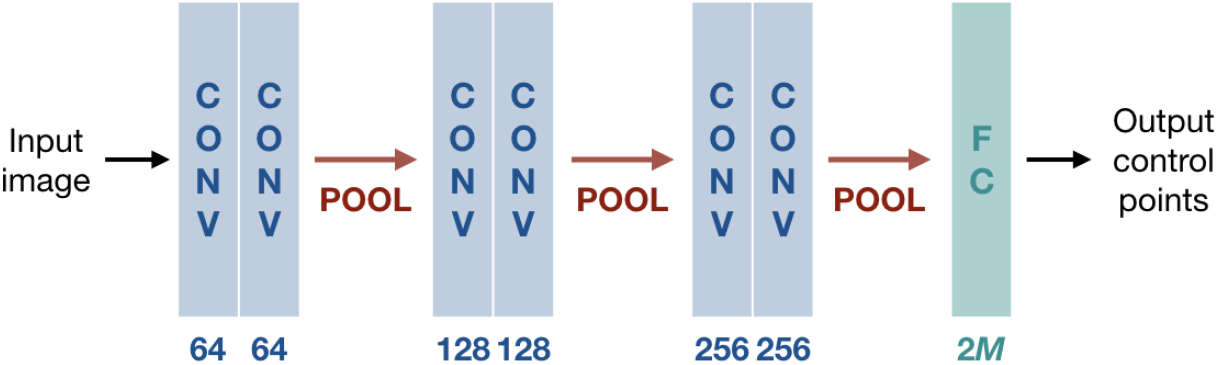
CNN Architecture. CONV: 3 ×3 leaky ReLU convolutional layer; POOL: 2 ×2 max-pooling layer; FC: fully connected layer; *M*: number of control points. The number of feature maps is indicated under each block.

Our training pipeline is illustrated in Figure 2. The input image goes through the CNN, which predicts the locations of the *M* control points {**c**[*k*]}_*k*=0,…,*M*−1_ of a spline model of the form (1), where the value of *M* and the spline generator *φ* have been chosen prior to training. The loss is then calculated between the discrete ground truth contour, extracted from the ground truth segmentation mask, and a sampled version of the predicted continuously-defined parametric curve. The network is updated accordingly for a given number of epochs using the ADAM optimizer [16].

**Fig. 2:**
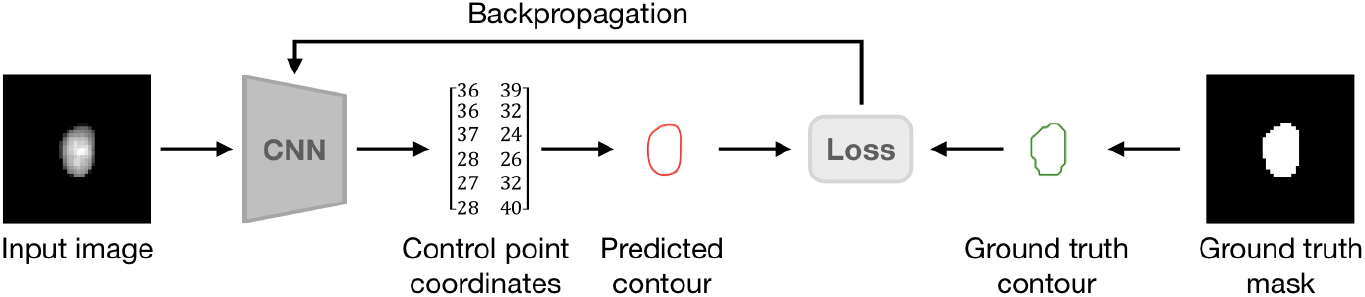
Training Pipeline. The input image goes through a CNN, which predicts the *M* control points {**c**[*k*]}_*k*=0,…,*M*−1_ of the parametric curve model (1). The loss reflects the discrepency between the ground truth (pixel-based) contour and the discretization of the predicted (continuous) one.

### 2.3 Loss Function

Objects in a segmented image are usually represented by a segmentation mask. To train our network, we extract the collections of pixels composing the contour of a ground truth segmentation mask using Satoshi and Abe’s algorithm [17] as implemented in OpenCV 3.4.2 [18]. The continuous curve the network predicts must faithfully capture the ground truth contour. Sampling the predicted continuous model, one should then retrieve the points located on the object contour (that is, the ground truth contour). We therefore define our loss as a Wasserstein (or Earth mover’s) distance [19] over ordered point sets.

We consider *A*, the set of *N* connected pixels composing the ground truth contour. By uniformly sampling *N* points on (1), we then retrieve *B*, an equally-sized set of points describing the predicted contour. Inspired by the work of [20] on image annotation, we train our network with the loss

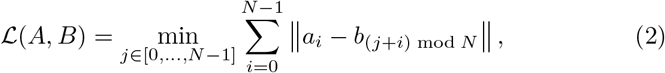

which corresponds to the minimum distance between *A* and all circular permutation of the elements in *B*. Classically, the Wasserstein distance is defined over unordered point clouds. Its calculation thus results in high computational complexity, and is in practice achieved relying on approximations. In our case, we can make use of the fact that successive points on the contour are inherently ordered, which dramatically reduces the complexity of the problem. Moreover, we can always ensure that *A* and *B* in (2) are equally-sized thanks to the continuously-defined nature of the predicted contour.

## 3 Experimental Results

In order to test the validity of our method, we consider the Kaggle 2018 Data Science Bowl dataset (also referred to as BBBC038v1) available from the Broad Bioimage Benchmark Collection [10]. It is composed of a diverse collection of images of cell nuclei, which aims at reflecting the type of images collected by research biologists at universities, biotechs, and hospitals. Nuclei in images vary in four different ways: the organism they are derived from, including but not limited to humans, mice, and flies; the way they have been treated and imaged, in terms of types of staining, magnification, and illumination; the context in which they appear, including cultured mono-layers, tissues, and embryos; and their physiological state, such as cell divison, genotoxic stress, and differentiation. The dataset faithfully reflects the variability of object appearance and image types in bioimages and is designed to challenge the generalization capabilities of a method across these variations We relied on the pre-defined stage1_train set, composed of 670 images, to train our network and perform cross-validation. We saved the stage1_test set, which consists of 65 images, to assess the performance of the network after training. Images are generally composed of more than one nucleus, but ground truth binary masks of each individual nuclei instance are provided for both sets. As we focus on continuously modelling the contour of a single object at once, we tile images according to bounding boxes around each individual nuclei. This step can be performed relying on a separate neural network trained for object detection, such as [21]. Ultimately, our final cross-validation and test sets are composed of respectively 29461 and 4152 tiles of isolated nuclei.

We divide the cross-validation set into 10 random partitions and follow a 10-fold cross-validation strategy. In each fold, we pick 9 partitions of the crossvalidation set for training and use the remaining partition as validation set to monitor performance. We ensure that each sample in the cross-validation set is used for training in 9 independent folds and for validation in the remaining one. We train the network for 100 epochs, keeping the learning rate as 0.0001. We record the evolution of the loss in training and validation to ensure convergence and monitor for overfitting. For each fold, we compute the median Dice score over the whole test set. The Dice score [22] is defined as Dice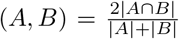, where *A* is the original ground truth mask and *B* the mask derived from the predicted contour. We obtain a median Dice score of 0.9562 ± 0.0014 over the ten folds, and provide visual illustration of high, intermediate, and low Dice score results in Figure 3.

**Fig. 3:**
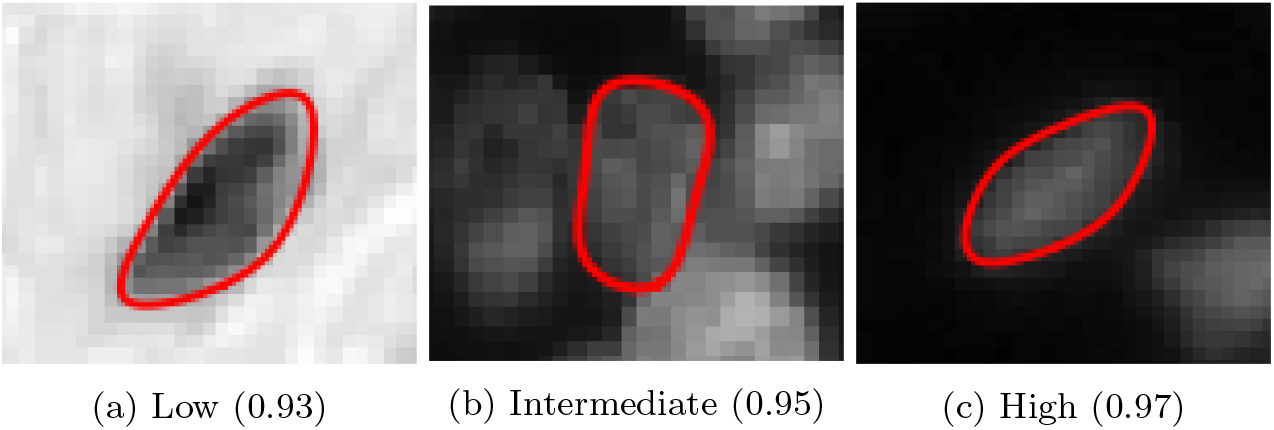
Predicted continuously-defined contours. We report examples of high quality (0.97), intermediate quality (0.95) and low quality (0.93) results from the BBBC038v1 stage1_test set after training on the BBBC038v1 stage1_train set.

Finally, in order to challenge the learning abilities of our rather shallow network architecture, we carry out a data ablation experiment. We keep the stage1 test set untouched and randomly ablate a fraction of the stage1_train set before training and carrying out 10-fold cross-validation. The depth of a network affects its ability to learn an appropriate data representation, but also dictates the amount of free parameters (weights) to be set. As we designed ours to offer a good trade-off between simplicity and performance, we here investigate the effect of training set size on prediction quality. In boxplots shown in Figure 4, we report the distribution of median Dice scores across folds on the full BBBC038v1 stage1_train set, as well as on ablated versions of it. No statistically significant decrease in performance is observed when ablating up to 90% of the stage1_train set. Prediction accuracy is clearly affected when training only on 10% of the data, but the median Dice nevertheless remains around 0.948. We provide source code to reproduce these experiments at gitlab.ebi.ac.uk/smandal/cpnet.

**Fig. 4:**
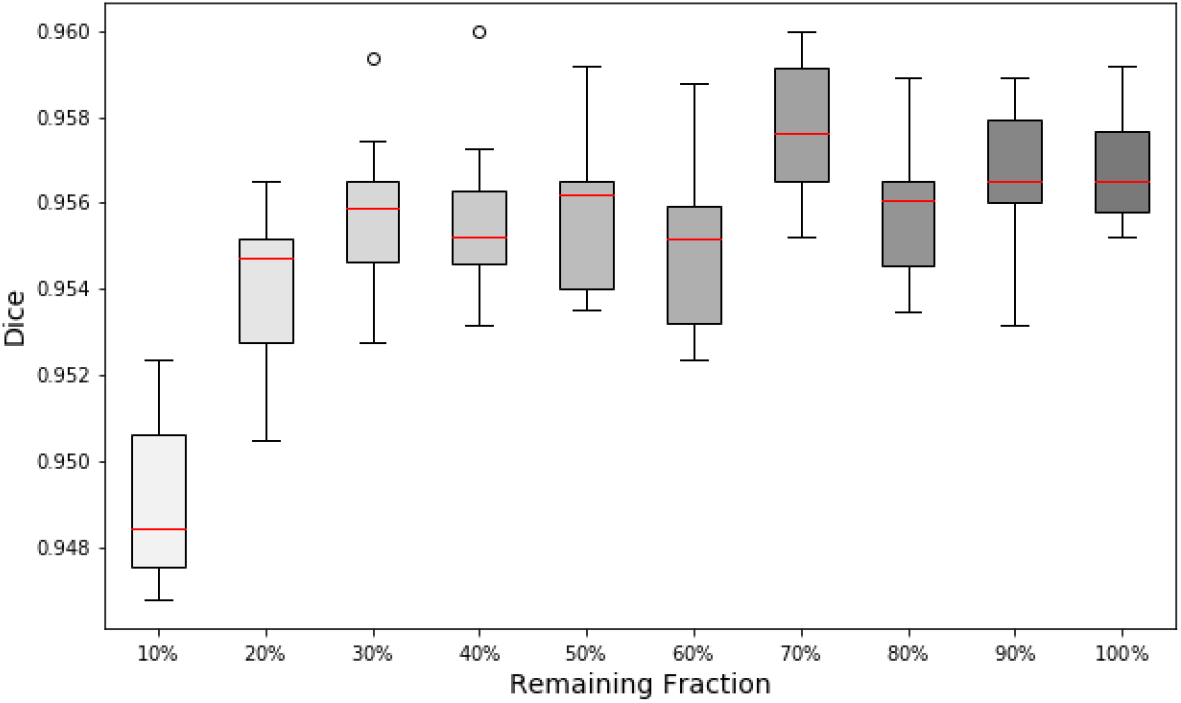
Data ablation study. Distribution of median Dice score across folds on the stage1_test set, when training on truncated versions of the stage1_train set.

## 4 Discussion

In this work, we propose a CNN-based pipeline to predict a continuous contour representation for objects in bioimages. The output of the network is a collection of discrete points that fully determine a continuously-defined parametric spline curve. Our method successfully allows to learn and predict continuous models of nuclei contours in images from the Kaggle 2018 Data Science Bowl dataset, which realistically reflects variations in the visual appearance of objects in bioimages. Here, the number of control points *M* and the basis function *φ* in (1) were determined prior to training and kept fixed. A natural future direction for this work aims at integrating these variables in the network and learning them from the nature of the objects to be represented. An additional worthy avenue would be to explore the generalization capabilities of the network on datasets featuring more complex biological object shapes.

## Acknowledgements

The authors are grateful to Dr James Klatzow for valuable discussions and helpful comments on the manuscript. This work is supported by EMBL core funding.

